# Gene-level structure of prior knowledge and perturbation responses informs target-level predictability in prior-informed models

**DOI:** 10.64898/2026.11.08.765139

**Authors:** Junha Park, Hyungtai Sim, Byungjin Hwang

## Abstract

**Background:** Predictive models of transcriptional responses to unseen genetic perturbations increasingly incorporate biological prior knowledge. However, it remains unclear how well prior knowledge reflects each target’s perturbation-response relationships and how closely prior-derived measures track target-level predictive performance. Here, we develop a gene-similarity framework to quantify agreement among eight sources of biological prior knowledge and between prior knowledge and co-function or co-regulation relationships derived from six Perturb-seq datasets.

**Results:** In genome-wide and essential-gene screens, prior agreement accounted for 30–47% of the variance in co-function agreement and 35–43% in co-regulation agreement. Agreement with Perturb-seq responses was higher for essential genes, lower for transcription factor and signaling genes, and varied with gene expression characteristics and cellular context. We compared prior agreement and prior-derived train–test similarity with measures based on the observed test response in tracking target-level predictive performance. Across three prior-informed models, predictions were more accurate for targets whose observed perturbation-response relationships better matched prior knowledge and that were more similar to training perturbations. Notably, prior agreement and prior-derived train–test similarity, calculated without the test response, tracked relative differences in target-level predictive performance. The strength of these associations varied across models and datasets and, for train–test similarity, across prior sources.

**Conclusion:** Our framework reveals gene- and context-level heterogeneity in how biological knowledge aligns with perturbation responses, and provides an a priori basis for assessing relative perturbation predictability before perturbation responses are measured.

## Background

Perturb-seq couples genetic perturbations with transcriptional readouts in single cells, enabling precise measurement of how perturbations reshape cellular states. Comprehensive mapping of genetic perturbations to rich transcriptional phenotypes under diverse cellular contexts has facilitated *in silico* prediction of perturbation responses, in which predictive models are trained to forecast transcriptional outcomes of unseen genetic perturbations. Recent predictive approaches increasingly incorporate biological prior knowledge, including co-expression patterns and Gene Ontology graphs [1–3], to enhance generalizability. Integrating multiple priors, such as pathway annotations and pretrained gene embeddings, can further improve prediction of unseen perturbation response and cross-dataset generalization [4,5].

However, prior knowledge is not equally informative for every gene or cellular context. Perturb-seq measures response in a specific cell type and experimental condition, whereas prior sources combine information from heterogeneous tissues, assays, annotations and observational datasets [5]. A gene may therefore be well characterized in existing databases but still show a context-specific perturbation response that those sources do not capture. Aggregate benchmark scores can obscure differences among individual targets. At the aggregate level, Systema showed that response components shared across perturbations can affect aggregate predictive performance [6].

Recent studies have also examined factors associated with target-level performance. GEARS examined knowledge-graph connectivity and model-derived uncertainty in relation to generalization performance [1], whereas TxPert related target knowledge and graph degree to perturbation-level performance [7]. PRESCRIBE further used target-to-training similarity and training-data quality to quantify model uncertainty and estimate confidence for unseen perturbations [8]. PRESAGE also related prediction performance to neighborhood density in aggregated prior embeddings [5]. These studies suggest that predictive performance for individual targets can be related both to prior information and to their relationship with the training set. Agreement among priors can be measured without the test response, but it does not by itself show how well those priors capture the target’s perturbation-response relationships. It therefore remains unclear how closely agreement among priors reflects agreement with each target’s perturbation-response relationships, and how prior-derived measures compare with measures based on the observed test response in tracking target-level predictive performance across models.

To address these questions, we considered agreement among prior sources and train–test similarity calculated from prior knowledge. We compared these prior-derived measures with agreement between prior knowledge and the target’s observed perturbation-response relationships, and with train–test similarity calculated from those responses. Within each model and dataset, we assessed how closely each measure tracked relative differences in predictive performance across held-out targets. Because genome-wide and essential-gene screens sample different target spaces and perturbation responses depend on cellular context, we also examined how these relationships varied across target sets and cellular contexts.

Accordingly, we developed a gene-similarity framework that systematically compares eight heterogeneous prior knowledge sources with six Perturb-seq datasets. We quantified how consistently each gene was represented across priors, which we termed prior agreement, and how well the priors matched two Perturb-seq-derived relationships: similarity of perturbation effects (co-function) and similarity of measured-gene responses across perturbations (co-regulation). Prior agreement accounted for substantial gene-to-gene variation in co-function and co-regulation agreement across genome-wide and essential-wide screens. These agreements also varied with perturbation-target annotations, gene expression characteristics, and cellular context. Even within the spliceosome, some genes showed consistently high co-function agreement across contexts, whereas others showed marked context-specific variation.

Across three prior-informed models, predictions were more accurate when prior knowledge better matched the target’s observed perturbation response and when the training set contained perturbations with similar effects. Prior agreement and prior-derived train–test similarity, calculated without the measured test response, were informative about relative predictive performance, although the strength of these associations varied across models and datasets.

By linking agreement among priors, agreement with observed perturbation-response relationships, and target-level predictive performance, our framework provides an a priori basis for comparing relative predictability across targets before the test response is measured. We envision using this framework to inform model- and context-aware prioritization of perturbations for experimental characterization.

## Results

### A gene-similarity framework linking prior knowledge to Perturb-seq responses and predictive performance

To enable direct comparison across heterogeneous prior knowledge sources and Perturb-seq datasets, we converted each source and dataset into gene-wise similarity profiles (**Fig. 1A**). Each gene was represented by a vector of its similarity to all other genes. In Perturb-seq, similarity between perturbation effects is referred to as co-function, whereas similarity between the responses of measured genes across perturbations is termed co-regulation.

**Fig. 1.**
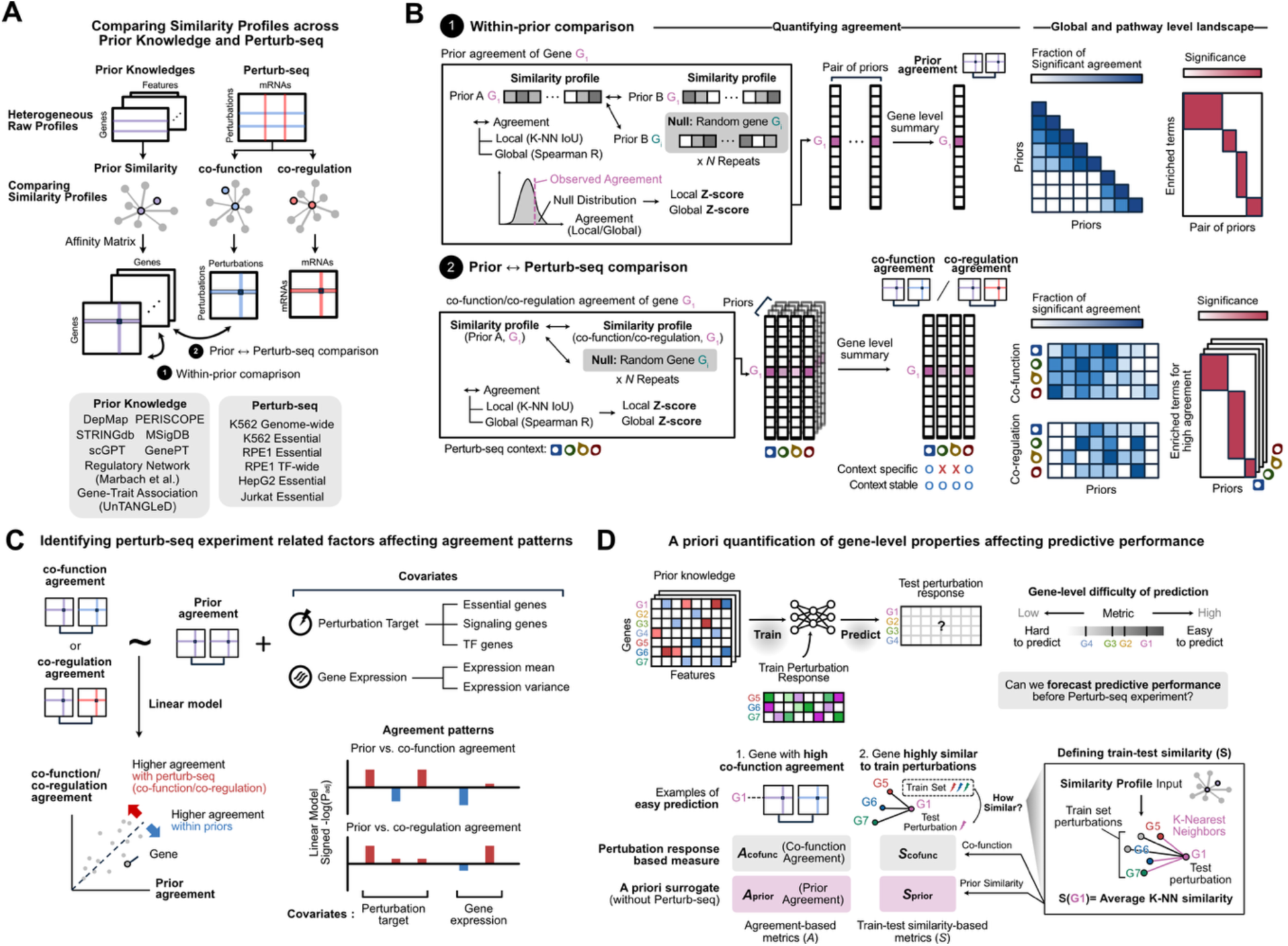
Overview of the study. (**A**) Gene-wise similarity profiles provide a common representation for comparing heterogeneous prior knowledge sources and Perturb-seq-derived response profiles. (**B**) Scheme for quantifying gene-level agreement within priors and between Perturb-seq data and priors. In each comparison, local and global agreement were calculated at the gene level. (**C**) Agreement metrics are used to assess associations with perturbation-target annotations and gene expression characteristics across cellular contexts. (**D**) Prior-derived measures of target-level prediction difficulty, calculated without the held-out response, are compared with response-informed measures that use the observed test response.

Using these shared representations, we compared similarity profiles at the gene level using the local overlap of nearest neighbors and global correlation across the full profile (**Fig. 1B**). These local and global measures were used for two types of comparison. First, comparisons among priors measured how consistently each gene was represented across prior resources. We summarized this consistency as prior agreement at the gene level. Second, comparisons between each prior and Perturb-seq measured how closely prior knowledge matched co-function or co-regulation relationships. These comparisons were summarized as co-function and co-regulation agreement. We then assessed the mean and standard deviation of co-function agreement across four essential-wide screens to distinguish context-stable from context-specific agreement patterns (**Methods**).

Next, we assessed how much variation in co-function and co-regulation agreement was explained by prior agreement, and whether perturbation-target annotations and gene expression characteristics explained additional variation. Using linear models, we estimated the associations of prior agreement and these additional factors with co-function or co-regulation agreement across cell lines (**Fig. 1C**).

Finally, we evaluated predictive performance for each held-out perturbation target, whose measured responses were excluded from model training and used only for evaluation, and related it to the gene-level relationship structures. Two relationships were considered: agreement-based metrics (*A*) and train–test similarity metrics (*S*). We asked whether prior agreement (*A*_prior_) and prior-derived train–test similarity (*S*_prior_), calculated without the test response, tracked relative predictive difficulty, and compared them with corresponding response-informed metrics. These assessed whether predictions were more accurate when prior knowledge better matched the similarity of the target’s perturbation effects to those of other targets (*A*_cofunc_), and when the training set contained perturbations with similar effects (*S*_cofunc_; **Fig. 1D**).

### Similarity profiles enable direct comparison of heterogeneous prior knowledge and Perturb-seq data

First, we collected eight prior knowledge sources spanning four types of biological information: graph-derived databases (MSigDB and STRINGdb), model-derived embeddings (GenePT and scGPT), CRISPR-based resources (DepMap and PERISCOPE), and genome-scale observations (regulatory networks from Marbach et al. and gene–trait associations from UnTANGLeD; **Fig. 2A**) [9–14]. To compare all eight priors using the same set of genes, we defined the 10,875 genes shared across all eight sources as the reference gene set. We then asked how much of each Perturb-seq dataset remained represented in this common gene space. Individual priors covered 75–100% of perturbation-targets and 86–99% of measured mRNAs, whereas the reference gene set covered 69–83% and 77–81%, respectively (**Fig. 2B**). Although requiring coverage by all eight priors reduced coverage relative to individual sources, the common gene space retained most perturbation targets and measured mRNAs across the Perturb-seq datasets.

**Fig. 2.**
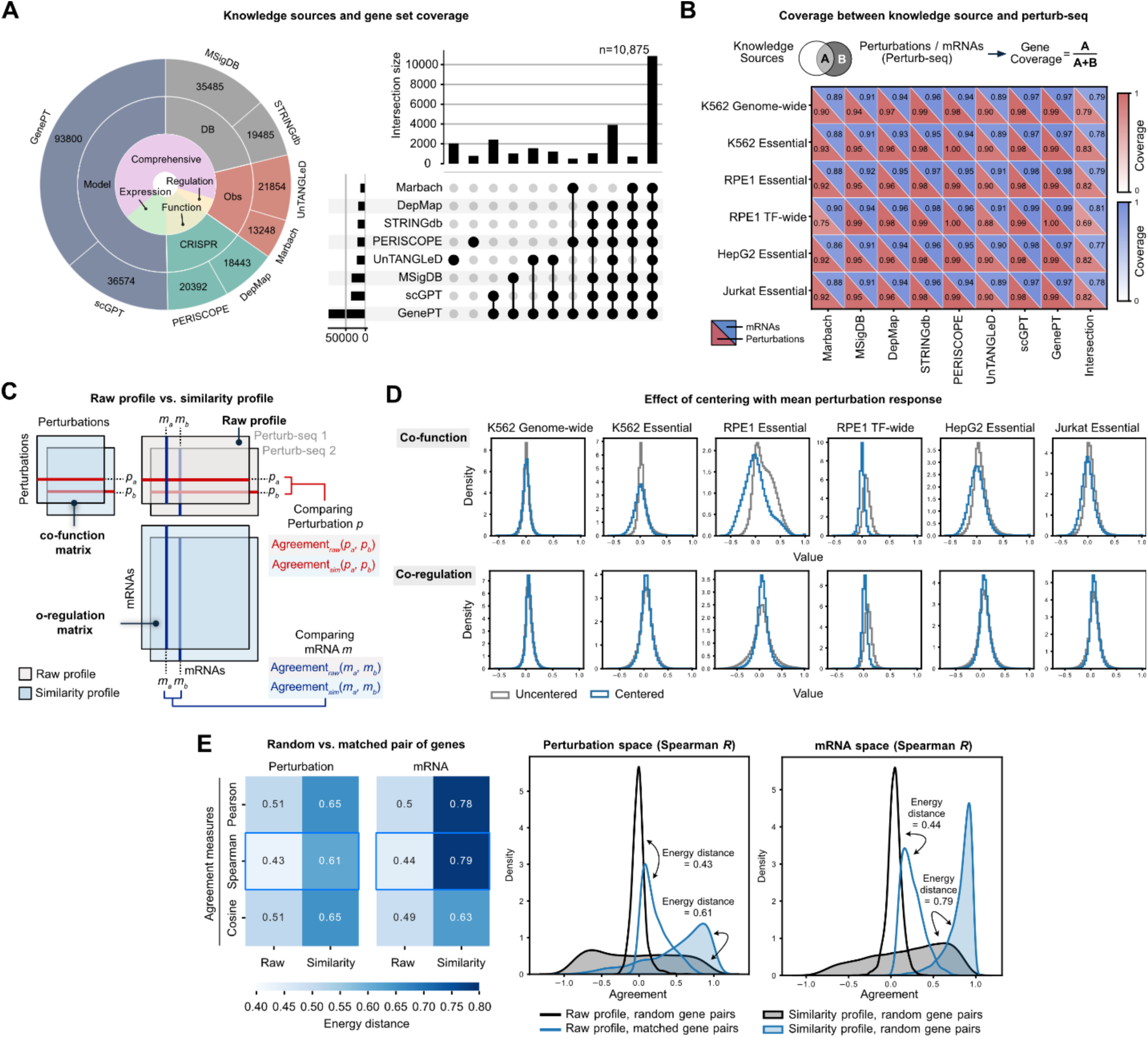
Gene–gene similarity profiles unify comparisons between Perturb-seq datasets and diverse biological knowledge sources. (**A**) Overview of the biological knowledge sources used for comparison. The sunburst plot summarizes knowledge-source categories and the number of genes covered by each source; the UpSet plot shows gene coverage shared across knowledge sources and Perturb-seq datasets. (**B**) Coverage of perturbation-targets and measured mRNAs in Perturb-seq datasets across knowledge sources. For each Perturb-seq– knowledge-source intersection, the upper triangle shows the fraction of measured mRNAs covered, and the lower triangle shows the fraction of perturbation-targets covered. Shading indicates coverage proportion. (**C**) Schematic of raw and similarity profiles in Perturb-seq data. Similarity between perturbations defines the co-function matrix, whereas similarity between mRNAs defines the co-regulation matrix. (**D**) Effect of mean-centering perturbation responses on pairwise similarity distributions. Gray outlines show uncentered distributions, and blue histograms show distributions after centering by the mean perturbation response. (**E**) Comparison of raw and similarity profiles using matched Perturb-seq datasets. K562 genome-wide data were restricted to the perturbation and mRNA spaces of the matched essential-wide screen. Heatmaps show median energy distances between matched- and random-gene-pair distributions across 100 repetitions and multiple similarity metrics. Density plots show representative distributions of Spearman correlations for raw and similarity profiles in the perturbation and mRNA spaces.

The prior resources and Perturb-seq datasets contain features with different dimensions and meanings, making their raw gene profiles difficult to compare directly. We therefore represented each gene by a similarity profile describing its relationship to all other genes (**Methods**). In Perturb-seq, similarity between perturbation profiles defines co-function, whereas similarity between measured-gene responses across perturbations defines co-regulation (**Fig. 1A and 2C**).

A response component shared across many perturbations can increase pairwise similarity through systematic variation rather than perturbation-specific effects [6]. Consistent with this, uncentered co-function values were shifted upward in all essential-gene and TF screens and for co-regulation in the RPE1 essential-wide dataset (**Fig. 2D**). We therefore mean-centered the response profiles before calculating similarity, removing the response component shared across perturbations. The resulting co-function and co-regulation distributions using all perturbation targets or measured mRNAs were comparable to those using only genes in the reference gene set (**Supplementary Fig. 1**).

We next tested whether similarity profiles improved cross-dataset comparability in Perturb-seq, where raw and similarity representations could be benchmarked directly. For a matched analysis, we restricted the perturbation and mRNA spaces of the K562 genome-wide screen to those of the K562 essential-wide screen. Across comparisons using Pearson correlation, Spearman correlation, and cosine similarity, similarity profiles more clearly separated matched gene pairs from randomly paired genes. Accordingly, the median energy distance increased from 0.43–0.51 to 0.61–0.65 in the perturbation space and from 0.44–0.50 to 0.63–0.79 in the mRNA space (**Fig. 2E**). These results show that similarity profiles improve cross-dataset discrimination of corresponding genes on both axes of the matched Perturb-seq response matrices, supporting their use in the following prior–prior and prior–Perturb-seq comparisons.

### Gene-level agreement reveals shared and distinct structure across prior knowledge sources

To assess how consistently each gene was represented across prior sources, we quantified both local and global agreement between its similarity profiles. Local agreement measured whether two sources identified the same closely related genes, whereas global agreement measured concordance across all gene relationships. Both measures were standardized against randomly paired genes to obtain Z-scores (**Methods; Supplementary Fig. 2A, B**). Local and global Z-score distributions were shifted above zero, indicating greater agreement between priors than expected by chance (**Fig. 3A**). The two scores were moderately correlated (Pearson’s *R* = 0.39; Spearman’s *ρ* = 0.33; **Supplementary Fig. 2C**). To summarize local and global agreement in a single score, we averaged them to define the gene agreement score (GAS), a combined measure of gene-level agreement for a given pair of similarity matrices.

**Fig. 3.**
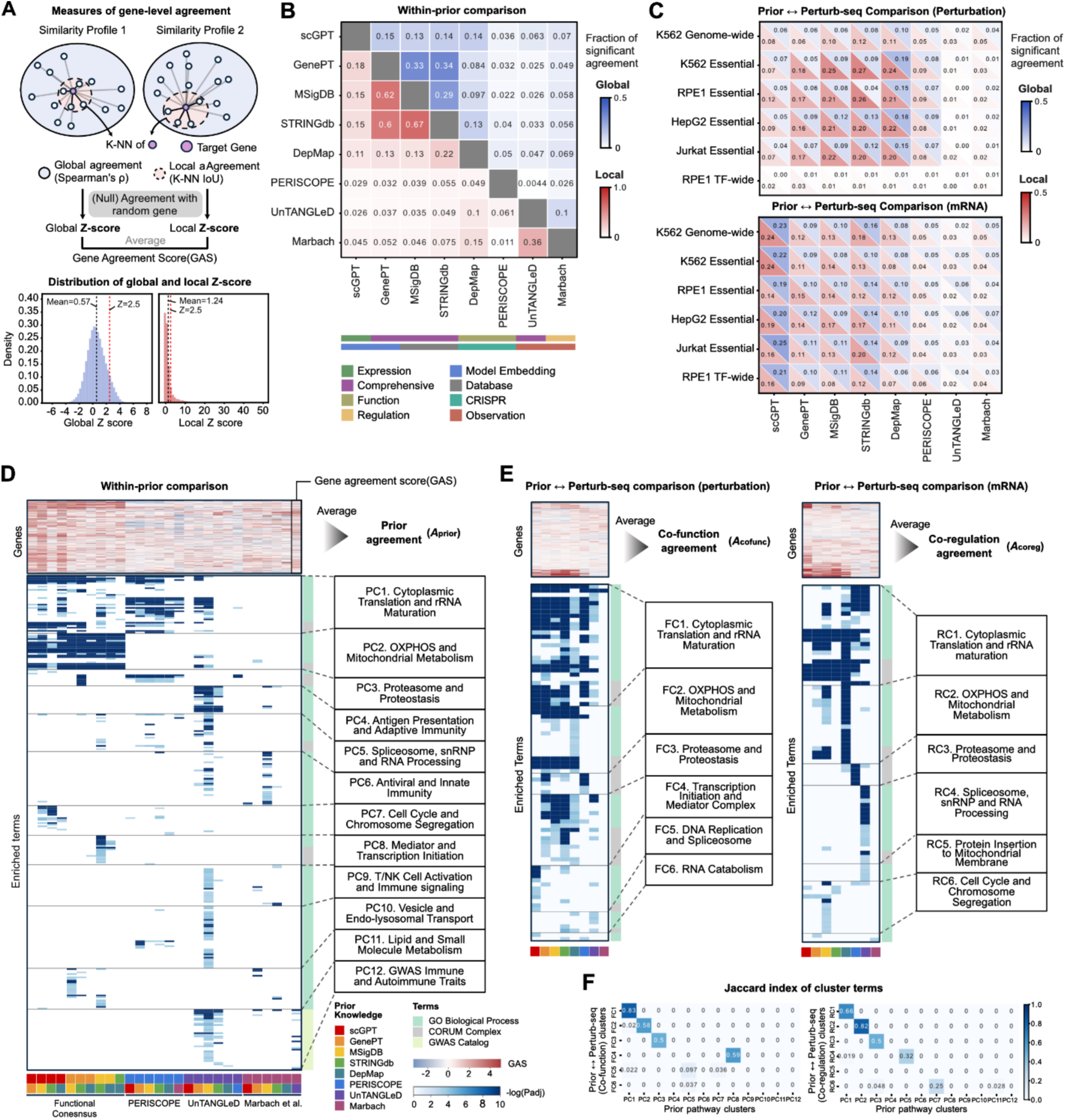
Gene-level agreement reveals shared and distinct organization across prior knowledge and perturbation responses. (**A**) Framework for quantifying gene-level agreement. Local agreement was measured by the intersection over union (IoU) of K-nearest-neighbor sets, and global agreement by the Spearman correlation between similarity profiles. Local and global agreement were standardized against random gene-pair null distributions, and their average was defined as the gene agreement score (GAS). The bottom density plot shows the distribution of standardized local and global Z-scores. (**B**) Fractions of genes with significant global (upper triangle, blue) or local (lower triangle, red) agreement between pairs of prior knowledge sources. Annotations below indicate prior categories. (**C**) Fractions of genes with significant agreement between each prior and perturbation-space co-function profiles (top) or mRNA-space co-regulation profiles (bottom) across six Perturb-seq datasets. Upper blue and lower red triangles denote global and local agreement, respectively. (**D**) Gene-wise GAS across 28 prior pairs (top) and gene set enrichment analysis of GAS (bottom). The average across prior pairs defines prior agreement, and enriched terms are grouped into 12 prior pathway clusters (PC1–PC12). (**E**) Corresponding analyses of co-function and co-regulation comparisons in the K562 genome-wide screen. Enriched terms are grouped into six co-function pathway clusters (FC1–FC6) and six co-regulation pathway clusters (RC1–RC6). Averages across eight priors define co-function and co-regulation agreement. (**F**) Jaccard index of enriched terms between prior pathway clusters and co-function or co-regulation pathway clusters.

For each prior pair, we next calculated the fraction of genes with significant local or global agreement above the threshold (Z-score > 2.5; **Fig. 3B**). Five sources—scGPT, GenePT, MSigDB, STRINGdb, and DepMap—showed consistently higher mutual agreement. The STRINGdb–GenePT pair showed local agreement above this threshold for 67% of genes. We refer to these five sources as the functional consensus group in subsequent analyses.

PERISCOPE, UnTANGLeD, and Marbach et al. were less concordant with the gene relationships shared by the five-prior group. This difference may reflect the distinct biological relationships and assays represented by these sources.

### Perturb-seq comparisons reveal context-specific agreement with priors

We next asked whether gene relationships represented in the priors were also reflected in co-function and co-regulation matrices from six Perturb-seq datasets (**Fig. 3C**) [15–18]. Gene-wise GAS patterns were highly concordant when the same priors were compared with the matched K562 essential-wide and genome-wide screens (**Supplementary Fig. 2D**), indicating that relative gene-level agreement patterns were preserved across the two datasets.

In K562, the essential-wide screen showed higher co-function agreement overall than the genome-wide screen (**Fig. 3C; Supplementary Fig. 2E**). In RPE1 cells, the TF-wide screen showed lower co-function agreement than the essential-wide screen. This contrast suggests that response to transcription factor perturbation measured in RPE1 cells may have distinct relationships beyond those inferred from prior similarity, consistent with the context dependence of transcription factor function [19,20]. In the matched K562 comparison, relative gene-wise agreement patterns were preserved despite differences in overall agreement; the RPE1 comparison further shows that agreement differs across perturbation target spaces within the same cell line.

The broad grouping observed in prior–prior comparisons was also evident in the prior–Perturb-seq comparisons. Within the functional consensus group, GenePT, MSigDB, STRINGdb, and DepMap consistently showed stronger agreement with co-function than scGPT across the six datasets, whereas scGPT showed the highest agreement with co-regulation. Thus, even priors with high mutual agreement differed in how they aligned with co-function and co-regulation. The stronger co-regulation agreement of scGPT is consistent with its pretraining on single-cell expression data and with the reported association between gene co-expression and perturbation-derived co-regulation [21]. In contrast, GenePT, MSigDB, STRINGdb, and DepMap more closely matched co-function relationships between perturbation targets. This difference between co-function and co-regulation is also consistent with previous benchmark results: scGPT-derived embeddings of mRNAs well reconstructed the transcriptional response to perturbations [22], whereas STRINGdb-derived embeddings of perturbations showed utility in extrapolating perturbation function [23].

### Pathway-level agreement identifies shared and comparison-specific biological programs

We next utilized pathway-level enrichment analysis to identify which biological programs contributed to the agreement patterns. For all 28 prior pairs, we performed gene set enrichment analysis (GSEA) on GAS using three complementary annotations spanning biological processes (GO Biological Process), protein complexes (CORUM), and disease-or trait-associated genes (GWAS Catalog) [24–26]. We organized enriched terms into 12 prior pathway clusters (PCs) shared across or specific to prior combinations (**Fig. 3D, Supplementary Table 1**). PC1 (Cytoplasmic translation and rRNA maturation) and PC2 (OXPHOS and Mitochondrial metabolism) were enriched mainly in comparisons within the functional consensus group, suggesting that these housekeeping programs are broadly represented across this group of prior resources. PC4 (Antigen presentation and adaptive immunity) and PC12 (GWAS immune and autoimmune traits) were enriched in pairs involving UnTANGLeD, consistent with its representation of gene-trait relationships from complex-trait data.

We applied the same analysis to GAS from each prior–Perturb-seq comparison in the K562 genome-wide screen and identified six co-function pathway clusters (FCs) and six co-regulation pathway clusters (RCs) (**Fig. 3E, Supplementary Table 2, 3**). Enriched terms for cytoplasmic translation and rRNA maturation, oxidative phosphorylation (OXPHOS) and mitochondrial metabolism, and proteasome/proteostasis overlapped with the PCs (**Fig. 3F**). Although scGPT showed lower overall co-function agreement in Fig. 3C, FC1 was enriched in the scGPT–co-function comparison, indicating pathway-specific agreement for translation and rRNA maturation genes. This is consistent with the reported performance of fine-tuned scGPT in predicting responses to perturbations of ribosomal genes [6].

Other programs differed between co-function and co-regulation. FC4 (transcription initiation and mediator complex) was specific to co-function comparisons, enriched for agreement with STRINGdb, MSigDB, DepMap, or GenePT. Conversely, RC6 (cell cycle and chromosome segregation) was specifically enriched for co-regulation comparison with MSigDB. Together, these patterns show that agreement among priors captures part, but not all, of the pathway structure observed in prior–Perturb-seq comparisons.

To quantify this relationship at the level of individual genes, we averaged GAS across the 28 prior pairs to define prior agreement and across the eight prior–Perturb-seq comparisons for each response matrix to define co-function and co-regulation agreement (**Fig. 3D, E**). We then asked how much prior agreement accounted for gene-to-gene variation in co-function and co-regulation agreement, and whether gene annotations and expression characteristics explained additional variation.

### Prior agreement accounts for substantial variation in co-function and co-regulation agreement

Using separate linear regression models, we tested how strongly prior agreement was associated with co-function and co-regulation agreement and whether perturbation-target annotations and gene expression characteristics explained additional variation (**Fig. 4A**). Target-gene annotations included essential genes from DepMap [15,16], genes associated with signaling pathways [27,28], and transcription factors (TFs; **Methods**) [18]. Expression-related covariates included the mean and variance of gene expression in control cells. We compared these associations across cell lines to assess their context dependence.

**Fig. 4.**
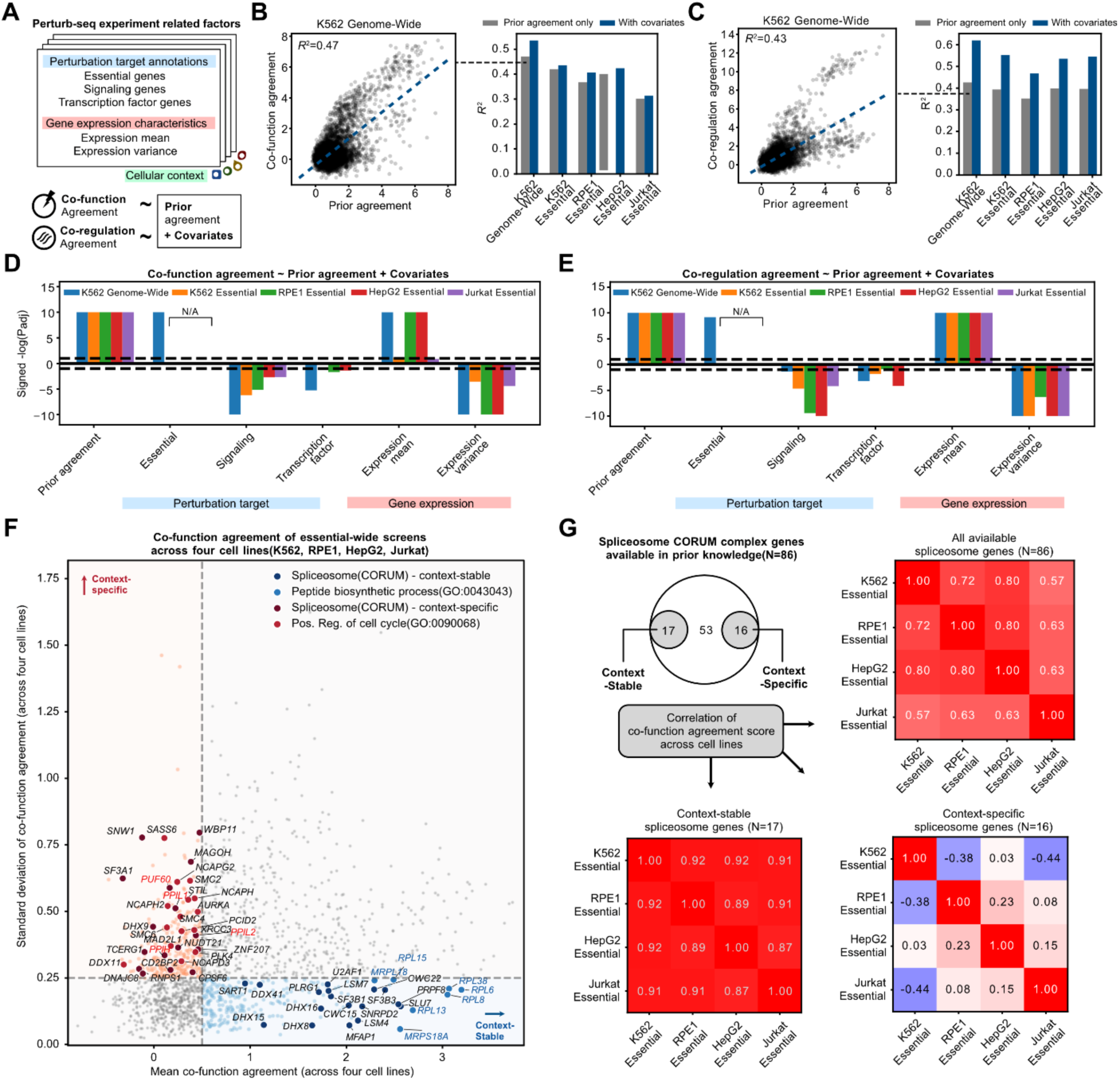
Gene-level correlates and context dependence of agreement between prior knowledge and perturbation responses. (**A**) Overview of the gene-level analysis. Co-function and co-regulation agreement were modeled using prior agreement and covariates related to perturbation-target annotations and gene expression characteristics, with analyses repeated across cellular contexts. (**B, C**) Association of prior agreement with co-function agreement (**B**) and co-regulation agreement (**C**) in the K562 genome-wide screen (left), and R^2^ for models containing prior agreement alone or prior agreement plus covariates across five Perturb-seq datasets (right). (**D, E**) Conditional associations of individual covariates with co-function (**D**) and co-regulation (**E**) agreement. Bars show sign(*β*)×−log10(FDR), and dashed lines indicate FDR = 0.05. Common-essential-gene status was estimable only in the K562 genome-wide screen. (**F**) Mean and standard deviation of co-function agreement for 1,625 perturbation-targets shared across the four essential-wide screens. Highlighted genes denote leading-edge members of selected enriched terms. (**G**) Pairwise Pearson correlations of co-function agreement across the four screens for all spliceosome genes represented in prior knowledge (n=86) and for context-stable (n=17) and context-specific (n=16) spliceosome subsets.

In the K562 genome-wide screen, prior agreement explained 47% of the variation in co-function agreement and 43% of the variation in co-regulation agreement (**Fig. 4B, C**). Across the four essential-wide screens, the corresponding values were 30–42% and 35–40%, respectively (**Supplementary Fig. 3A, B**). Genes represented more consistently across priors therefore tended to show stronger co-function and co-regulation agreement. Adding gene annotations and expression characteristics further increased the explained variance, although the increase was smaller than that explained by prior agreement alone (Δ*R*² = 0.011–0.064 for co-function agreement and 0.115–0.194 for co-regulation agreement). The additional contribution was larger for co-regulation than for co-function agreement.

The RPE1 TF-wide screen differed from the genome-wide and essential-wide screens: prior agreement explained only 1% of the variation in co-function agreement, compared with 29% for co-regulation agreement (**Supplementary Fig. 3A, B**). The corresponding RPE1 essential-wide screen did not show the same divergence, indicating that this pattern was not explained by cell line alone and may instead reflect properties of TF perturbations that depend on target and experimental context [19,20].

### Gene-level agreement varies with perturbation-target annotation, gene expression characteristics, and cellular context

We next examined whether perturbation-target annotation and expression characteristics were associated with co-function and co-regulation agreement after accounting for prior agreement (**Methods**). In the K562 genome-wide dataset, gene essentiality was positively associated with both co-function and co-regulation agreement after adjustment for prior agreement and the other covariates (**Fig. 4D, E; Supplementary Fig. 3C**). Together with the higher co-function agreement observed for essential genes in the K562 genome-wide screen (**Supplementary Fig. 2E**), this result suggests that perturbation responses of essential genes are more consistently represented in prior knowledge. Transcription factor genes were associated with lower co-function and co-regulation agreement in the K562 genome-wide dataset (**Fig. 4D, E; Supplementary Fig. 3D**). Signaling genes were also negatively associated with both agreement measures across the cell lines (**Fig. 4D, E; Supplementary Fig. 3E**). Higher mean expression and lower expression variance were generally associated with greater co-function and co-regulation agreement after adjustment for the other covariates.

The association of co-function agreement with prior agreement varied more across four essential-wide screens than did the corresponding association for co-regulation (**Supplementary Fig. 3A**). To examine this variation among comparable targets, we focused on co-function agreement for genes shared across the four essential-wide screens.

Across four essential-wide screens, we identified genes with consistently high or context-specific co-function agreement (**Fig. 4F**). We defined context-stable genes as those with high mean co-function agreement and low standard deviation, and context-specific genes as those with low mean and high standard deviation (**Methods; Supplementary Fig. 3F**). Pathway enrichment analysis showed consistently high agreement for peptide-biosynthesis and mRNA-processing programs (**Supplementary Fig. 3H**). Cell-cycle-regulatory programs showed greater context-dependent variation (**Supplementary Fig. 3G**), whereas spliceosome-related programs were enriched in both context-stable and context-specific analyses (**Supplementary Fig. 3G, H**).

This pattern suggested that context dependence may differ among spliceosome components. Co-function agreement was strongly correlated across cell lines for context-stable spliceosome genes but substantially less concordant for the context-specific subset, particularly in comparisons of K562 with RPE1 or Jurkat cells (**Fig. 4G**). Context-stable spliceosome genes included *DDX41,* which was previously reported to show highly correlated transcriptome-wide effects across cell lines [16]. Context-specific spliceosome genes included immunophilins such as *PPIL1*, *PPIL2*, and *PPIH* [29], and *PUF60*, which is associated with cell-cycle regulation [30]. Thus, prior–Perturb-seq agreement varies not only across broad gene categories and pathways but also among components of the same molecular complex, depending on cellular contexts.

### *A priori* assessment of target-level predictability in prior-informed predictive models

We then asked whether two prior-derived measures—(1) prior agreement and (2) train–test similarity—calculated without the observed test response were associated with predictive performance for individual held-out targets. We compared their forecastability with that of corresponding response-informed measures. Using a benchmark of three prior-informed models, GEARS, scLAMBDA and PRESAGE [1,5,31], on held-out perturbations [32], we quantified predictive performance for each test perturbation. To distinguish how well a metric tracks target-level performance from the model’s predictive accuracy, we defined forecastability as the Pearson correlation across test perturbations between a predefined predictability metric and observed predictive performance (**Fig. 5A**). Thus, a high positive forecastability value indicates that perturbations with higher metric values tend to have higher predictive performance. We compared forecastability across predictive models, perturbation-target sets, and cellular contexts.

**Fig. 5.**
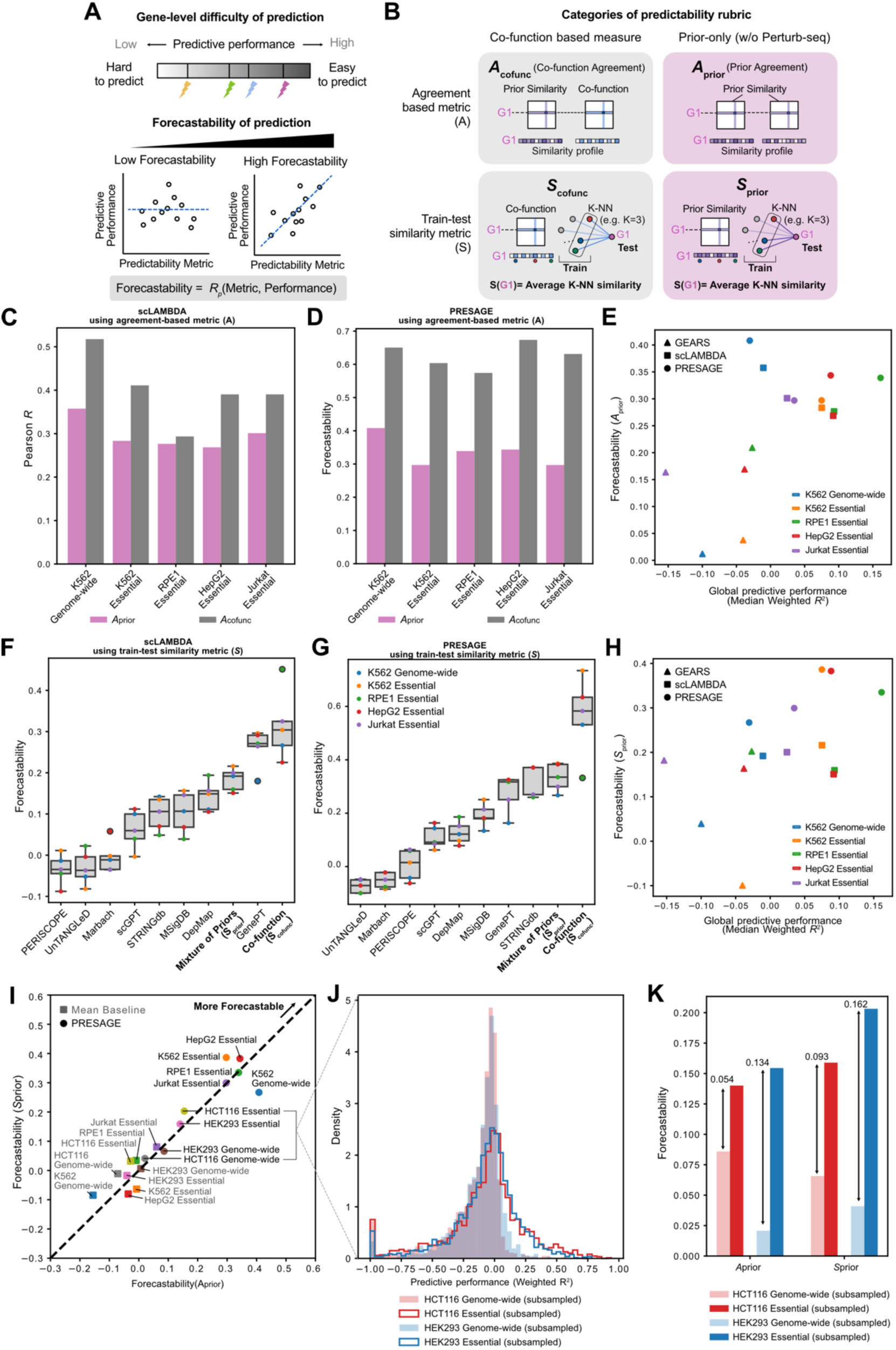
Forecastability of prior-informed predictive performance. (**A**) Schematic of forecastability, defined as the Pearson correlation across held-out targets between a predictability metric and observed predictive performance. (**B**) Categories of predictability metrics. Agreement-based (top) and train–test similarity (bottom) metrics were calculated as response-informed measures using the observed test response (left) or prior-derived measures without that response (right). (**C, D**) Forecastability of scLAMBDA (**C**) and PRESAGE (**D**) performance using agreement-based metrics (A). (**E**) Global predictive performance (median weighted *R*^2^ across all test perturbations) of GEARS, scLAMBDA, and PRESAGE across cell lines (x axis) and corresponding forecastability using prior agreement (*A*_prior_) (y axis). (**F, G**) Forecastability of scLAMBDA (**F**) and PRESAGE (**G**) performance using train–test similarity metrics (*S*). (**H**) Global predictive performance of GEARS, scLAMBDA and PRESAGE across cell lines (x axis) and corresponding forecastability using train–test similarity (y axis). (**I**) Forecastability of PRESAGE performance across nine combinations of cell lines and perturbation-target sets. (**J**) Distribution of PRESAGE performance (weighted *R*^2^ across test perturbations) for genome-wide and essential gene-restricted subsets of the HCT116 and HEK293 Perturb-seq experiments. Genome-wide screens were subsampled to match the number of training and test perturbations in the essential-wide restriction. (**K**) Forecastability of PRESAGE performance using *A*_prior_ or *S*_prior_ as a metric for genome-wide and essential gene-restricted subsets of the HCT116 and HEK293 Perturb-seq experiments.

We evaluated two types of metrics: agreement-based (*A*) and train–test similarity (*S*) metrics. Each type was calculated from prior knowledge without the test response or from the observed perturbation responses (**Fig. 5B**). This yielded a two-by-two comparison of the relationship measured (*A* or *S*) and the information used (prior-derived or response-informed). Agreement-based metrics compare gene relationships, whereas train–test similarity metrics relate each test target to perturbations represented in the training set.

For agreement-based metrics, co-function agreement (*A*_cofunc_) served as the response-informed metric, whereas prior agreement (*A*_prior_) served as its prior-derived counterpart. For train–test similarity metrics, we defined *S* within a given similarity matrix as the average similarity between each test perturbation and its *K* nearest training perturbations. Computing *S* from the co-function matrix yielded *S*_cofunc_, whereas computing *S* separately from each prior matrix yielded prior-specific *S* values. We averaged these prior-specific *S* values across the five priors in the functional consensus group to define *S*_prior_ (**Fig. 5B**). The *S* metrics were stable across the evaluated values of *K* (**Supplementary Fig. 4A; Methods**).

Across the five datasets, co-function agreement (*A*_cofunc_) showed higher forecastability than prior agreement (*A*_prior_) for all three models (**Fig. 5C, D; Supplementary Fig. 5A**). Thus, agreement between prior knowledge and the observed test-perturbation response tracked target-level predictive performance more closely than agreement among prior sources alone. Notably, *A*_prior_ showed positive forecastability for both PRESAGE (Pearson’s *R* = 0.297–0.408) and scLAMBDA (*R* = 0.268–0.358; **Fig. 5E**), indicating that prior knowledge alone provided information about relative predictive performance. Forecastability was weaker for GEARS (*R* = 0.012–0.209; **Fig. 5E**), with the large differences between GEARS and PRESAGE observed in the K562 genome-wide and essential-wide screen (**Fig. 5D; Supplementary Fig. 5A**).

To place these gene-level associations in a biological context, we compared PRESAGE predictive performance, *A*_prior_, and *A*_cofunc_ across functional gene sets in the K562 essential-wide screen (**Supplementary Fig. 5B**). The cytoplasmic ribosomal subunit sets combined relatively high co-function agreement with positive predictive performance. Other gene sets showed distinct relationships between prior agreement and performance: the Pol I and rRNA biosynthesis set showed higher predictive performance but lower *A*_prior_ than the mitochondrial 39S ribosomal subunit set.

Next, we examined train-test similarity. *S*_cofunc_ generally showed higher forecastability than *S* derived from individual prior sources (**Fig. 5F, G; Supplementary Fig. 5C**). Prior-derived *S* metrics that most closely tracked target-level predictive performance varied across models. In GEARS, scGPT-derived *S* showed higher forecastability than *S*_cofunc_ in most datasets.

For scLAMBDA, GenePT-derived S showed forecastability comparable to that of *S*_cofunc_ and higher than that of *S*_prior_ in four of the five datasets. In contrast, *S*_prior_ generally showed slightly higher forecastability than individual-prior *S* metrics for PRESAGE (**Fig. 5G**). The arithmetic mean of train–test similarity values across the five functional-consensus priors therefore did not always yield the highest observed correlation with target-level performance. *S*_prior_ showed higher forecastability for PRESAGE (*R* = 0.267–0.386) than for scLAMBDA (*R* = 0.151–0.216) or GEARS (*R* = −0.100–0.202; **Fig. 5H; Supplementary Fig. 5C**). This may reflect the differences in prior usage across evaluated prior-informed models. Together, these analyses show that prior agreement and prior-derived train–test similarity can track relative differences in predictive performance across held-out targets, but the strength of these associations varies across predictive models and, for train–test similarity, across prior sources.

### Forecastability varies across cellular contexts and perturbation-target sets

Because PRESAGE showed relatively high forecastability for both *A*_prior_ and *S*_prior_, we next examined cross-context variation within PRESAGE. We extended the PRESAGE benchmark to genome-wide Perturb-seq datasets from HEK293 and HCT116 cells [17]. For each dataset, we also constructed an essential-gene-restricted evaluation by limiting both training and test perturbations to essential genes.

Across nine combinations of cellular contexts and perturbation-target sets, forecastability based on prior agreement (*A*_prior_) and prior-derived train–test similarity (*S*_prior_) varied substantially (**Fig. 5I**). Both metrics showed higher forecastability for PRESAGE than for the mean-predictor baseline, which relies only on the overall statistics of training perturbation responses, across these settings. However, HEK293 and HCT116 showed lower forecastability than the other screens in both genome-wide and essential-gene-restricted analyses.

To assess whether forecastability differed between target sets of the same size, we matched the numbers of training and test perturbations between the genome-wide and essential-gene-restricted evaluations within each cell line (**Methods**). The essential-gene-restricted evaluations showed higher predictive performance and forecastability, for both *A*_prior_ and *S*_prior_ (Fig. 5J, K).

The metrics therefore tracked differences in target-level performance more closely in the essential-gene-restricted evaluations, even after matching perturbation counts. However, forecastability remained comparatively lower than in the other essential-wide screens, indicating that essential-gene restriction alone did not remove the differences between datasets.

### High-similarity training sets yield better predictions for fixed targets under matched prior agreement

To extend the assessment of target-level predictability to training-set selection, we asked whether prior similarity could guide the choice of training perturbations for specified test targets. In a retrospective training-set selection analysis, we compared high- and low-similarity training sets while holding the test targets fixed and matching the distributions of *A*_prior_ in the training sets (**Supplementary Fig. 6A**). Across five randomly selected test sets of 100 perturbations each, high-similarity training sets yielded higher predictive performance than low-similarity training sets at each evaluated training-set size (**Supplementary Fig. 6B**). These analyses show how prior-derived measures can inform both the assessment of relative target-level predictability and the selection of training perturbations for specified targets.

## Discussion

Prior-informed models such as GEARS, PRESAGE, and TxPert [1,5,7] incorporate biological knowledge to predict responses to unseen perturbations. Recent benchmarks have shown that systematic variation can affect aggregate predictive performance [6]. Comparisons with simple baselines further show that conclusions about model performance depend on the evaluation task and metric [22, 32]. In addition, differences between training and test perturbation sets can also influence overall benchmark performance [33]. However, aggregate scores can mask variation in predictive performance across individual perturbations. To address these gaps, we introduced a gene-level framework linking agreement among priors, agreement with observed perturbation-response relationships, and perturbation-specific predictability. Prior agreement and prior-derived train–test similarity, calculated without the test response, were informative about relative predictive performance across held-out targets, with associations varying across models and datasets.

Prior–response agreement was biological program-specific and differed between co-function and co-regulation. Agreement was concentrated in specific biological programs, with translation and rRNA maturation represented in both co-function and co-regulation comparisons and transcription initiation and the mediator complex specific to co-function comparisons. scGPT showed the strongest co-regulation agreement, whereas GenePT, MSigDB, STRINGdb, and DepMap more closely matched co-function.

At the gene level, we next examined gene-to-gene variation in co-function and co-regulation agreement. Prior agreement accounted for a substantial component of this variation, with additional variation associated with perturbation-target annotations and gene expression characteristics. Spliceosome genes illustrated that co-function agreement can differ across cellular contexts even among genes within the same molecular complex. Prior agreement therefore captured a substantial but incomplete component of gene-level prior–response agreement, with further variation associated with target annotations, expression characteristics, and cellular context.

Forecastability complements aggregate predictive performance by assessing whether gene-level measures distinguish relatively easier and harder perturbations to predict. For agreement-based metrics, co-function agreement showed higher forecastability than prior agreement across the three models (**Fig. 5C, D**; **Supplementary Fig. 5A**). Notably, prior agreement was positively correlated with target-level predictive performance in scLAMBDA and PRESAGE (Pearson’s *R* = 0.268–0.358 and 0.297–0.408, respectively). For train–test similarity, the metric based on observed perturbation responses generally showed higher forecastability than prior-derived metrics, but some metrics derived from individual priors showed comparable or higher forecastability in specific model–dataset settings. Thus, response-informed agreement metrics consistently showed higher forecastability than their prior-derived counterparts, whereas this advantage was less consistent for train–test similarity.

These differences suggest that prior-derived measures are not equally informative across predictive models. In PRESAGE, averaging train–test similarity across the functional consensus priors generally showed higher forecastability than metrics derived from individual priors. By contrast, specific priors were more informative in the other models: GenePT-derived similarity showed higher forecastability than the five-prior average in four of five datasets in scLAMBDA, whereas scGPT-derived similarity showed relatively high forecastability in GEARS (**Fig. 5F, G**; **Supplementary Fig. 5C**). The informativeness of prior-derived train–test similarity is therefore model-dependent rather than a fixed property of the prior itself, and averaging across priors does not necessarily yield the most informative train–test similarity measure for every model.

Taken together, these model-dependent associations suggest that prior-derived metrics may help distinguish easier and harder targets to predict within a given model and dataset. Because they do not require the test response, these metrics can be calculated before it is measured.

The extended PRESAGE analyses in HEK293 and HCT116 further showed higher forecastability when both training and test perturbations were restricted to essential genes, even after matching the number of target perturbation.

Because these metrics can be calculated without the test response, their associations with predictive performance can inform relative confidence in target predictions and help prioritize perturbations for experimental characterization. The fixed-target training-set analysis further extends their use to selecting training perturbations. High-similarity training sets yielded better predictions than low-similarity sets with matched distributions of *A*_prior_ (**Supplementary Fig. 6**).

This study has several limitations. First, both families of metrics were associated with predictive performance, yet substantial variation remained, suggesting that other factors, such as sequencing depth or cell numbers per perturbation, may also influence predictive performance. Differences across screens also cannot be attributed to cellular context alone because target composition and experimental protocols differ. Second, prior-derived estimates may be constrained by the incomplete coverage and context specificity of current prior knowledge.

Requiring coverage by all eight priors may favor well-characterized genes, while shared information and reuse of the same prior resources limit the independence of the agreement measures. Finally, our analyses were restricted to perturbation responses measured in the mRNA expression space. Whether similar agreement patterns and prior-derived associations with predictive performance extend to protein-level, multi-omic, or imaging-based phenotypes remains to be established [34–37].

## Conclusions

Our analyses show that the usefulness of prior knowledge depends on both the biological relationship of interest and the model in which the prior is used, and that these patterns vary across cellular contexts and perturbation targets. Agreement among priors explained part of the gene-to-gene variation in how well prior knowledge matched observed perturbation responses. Together with prior-derived train–test similarity, it also provided information about relative predictive performance across held-out targets before their responses were observed. These metrics could inform experimental prioritization strategies, including active-learning frameworks [38], by highlighting perturbations for which existing information may provide limited support for prediction. Large-scale Perturb-seq datasets spanning broader target sets and cellular contexts could help evaluate the generalizability of prior-derived measures of perturbation predictability. Prospective studies can test whether experimentally characterizing such perturbations improves subsequent prediction of unseen perturbations.

## Methods

### Prior knowledge sources

Eight knowledge sources were assembled as comparative priors: model-derived embeddings (GenePT and scGPT) [11,12]; graph-like resources (MSigDB embeddings from medium-confidence edges and STRINGdb embeddings) [9,10]; resources derived from observational studies (Marbach et al. and the UnTANGLeD gene–trait association matrix) [19,39]; and CRISPR knockout (KO) datasets (PERISCOPE and DepMap) [13,14]. For the GenePT embedding, graph-like resources, and DepMap dataset, we used preprocessed matrices generated by the PRESAGE study as gene-feature embeddings [5]. GenePT embeddings were corrected using the mean embedding across all genes in the reference gene set because random gene pairs showed high basal correlation. Source-specific preprocessing was performed for resources not directly adopted from PRESAGE. scGPT gene embeddings were extracted from the pretrained scGPT blood model by encoding gene-vocabulary IDs; the resulting gene-by-embedding matrix was used. We loaded the gene-feature matrix measured by PERISCOPE in A549 cells and converted the Marbach et al. regulatory network from chronic myelogenous leukemia (CML) cells into a gene-by-TF weight matrix, setting missing edges to zero. For UnTANGLeD gene–trait association data, we retrieved the MultiXcan gene–trait association matrix and mapped Ensembl gene identifiers to gene symbols. The resulting matrices were used to generate gene–gene similarity profiles.

### Perturb-seq datasets

Raw count matrices from Replogle et al. (2022), Nadig et al. (2025), and the Huang et al. (2025) datasets were retrieved according to the Data Availability instructions [15–17]. To prepare similarity matrices from Perturb-seq datasets, we excluded perturbation labels represented by fewer than 10 cells and retained expression features with a mean UMI count greater than 0.01 across the remaining cells. Counts were normalized for library size within each batch by scaling each cell to the batch-specific median read depth across retained features. For each perturbation, z-scores were averaged across all corresponding cells to generate an effect-size matrix, with or without feature mean-centering.

### Calculation of gene–gene similarity matrices

Gene–gene similarity matrices were calculated separately for each prior knowledge source and Perturb-seq dataset. For prior knowledge sources, similarity matrices were computed from the corresponding gene-feature matrices. Cosine similarity was used for PRESAGE-derived embeddings (GenePT, STRINGdb, MSigDB, and DepMap) and scGPT embeddings. For the other prior knowledge sources (PERISCOPE, Marbach et al., and UnTANGLeD), gene-feature matrices were converted to similarity matrices using Spearman correlation across features. For Perturb-seq datasets, similarity matrices were computed from perturbation-level effect-size matrices. Cosine similarity, after centering by the mean across all perturbations, was used to compare response profiles between perturbed genes.

### Estimation of gene-level agreement between similarity matrices

Gene-level agreement between two similarity matrices was quantified using local and global measures of their similarity profiles. For a given gene, its similarity profile was defined as the vector of similarity values between that gene and all other shared genes, excluding self-similarity.

Local agreement was measured as the intersection over union (IoU) between the top-K nearest-neighbor gene sets identified from the two similarity profiles. We selected K=100 for the local agreement to ensure sufficient neighborhood coverage and robustness (**Supplementary Fig. 2A and 2B**). Global agreement was measured as the Spearman correlation between the full similarity profiles from the two matrices. To estimate agreement expected by chance, we generated a null distribution by comparing randomly paired genes across the two matrices in 1,000 repetitions. The observed local IoU and global Spearman correlation were converted to local and global Z-scores using the mean and standard deviation of the corresponding null distribution.

A gene agreement score (GAS) was defined as the average of the local and global Z-scores for each gene–matrix pair. Genes with either a local or global Z-score > 2.5 were considered to show significant agreement. Prior agreement was calculated by averaging GAS across all pairwise comparisons among prior knowledge sources. Co-function and co-regulation agreement were calculated by averaging GAS across comparisons between each prior knowledge source and the corresponding Perturb-seq-derived co-function or co-regulation matrix, respectively.

### Gene set enrichment analysis

Gene set enrichment analysis (GSEA) was performed using *gseapy* [24–26,40] to evaluate the agreement of prior-wide and prior-specific signatures with GO Biological Process, CORUM complex, and GWAS Catalog gene sets. Among significant enrichment results (FDR < 0.001), clusters were annotated based on their biological roles.

### Estimation of covariate-adjusted alignment between agreement measures

For each Perturb-seq dataset, we fitted separate ordinary least-squares regression models with co-function or co-regulation agreement as the outcome and prior agreement as the predictor of interest, with one gene as the unit of analysis. Covariate-adjusted analyses included the K562 genome-wide screen and four essential-wide screens (K562, RPE1, HepG2, and Jurkat), retaining only genes with both prior-agreement and outcome values. Prior-agreement-only regressions were also summarized for the RPE1 TF-wide screen (**Supplementary Fig. 3A, B**). Full models included binary indicators of common-essential genes, signaling genes, and transcription factors, together with continuous covariates for gene-wise mean expression and expression variance. Common-essential, signaling, and transcription factor genes were defined using the DepMap CRISPRInferredCommonEssentials set; the union of genes assigned to KEGG 2026 canonical and system-specific signaling pathways in the immune, endocrine, circulatory, and nervous systems; and targets from the Southard et al. transcription factor screen [18,41], respectively. Mean expression and expression variance were calculated across control cells, with variance calculated as the population variance. Missing covariate values were mean-imputed. The common-essential indicator was included only in the K562 genome-wide analysis.

All models included an intercept, and the coefficient for prior agreement was reported as the covariate-adjusted alignment estimate. Additional variance explained by the covariates was quantified as 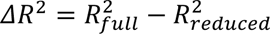, where the reduced model contained prior agreement alone.

Two-sided P values for regression coefficients were adjusted within each model using the Benjamini–Hochberg procedure, and associations were summarized as *sign* (*β*) × −*log* 10(*q*), where β is the regression coefficient and q is the adjusted P value. Variance inflation factors were calculated for the predictors in the full model.

### Identification of context-specific and context-stable agreement patterns

Co-function agreement patterns were classified as context-specific or context-stable by thresholding the mean and standard deviation of co-function agreement across four cell lines (K562, RPE1, HepG2, and Jurkat). Based on inspection of these distributions, GSEA was performed among genes with a standard deviation < 0.25, using mean co-function agreement as the gene-wise score, to identify pathways with context-stable agreement. Among genes with a mean < 0.5, GSEA was performed using the standard deviation of co-function agreement to identify pathways with context-specific agreement. Among enriched pathways (Benjamini– Hochberg-adjusted *P* < 0.1), the five with the highest normalized enrichment scores (NESs) were used to annotate gene sets harboring context-specific and context-stable agreement patterns.

### Calculation of forecastability metrics

Agreement-based (*A*) and train–test similarity (*S*) metrics were calculated as prior-derived or response-informed measures. *A*_prior_ compares priors with one another, whereas *A*_cofunc_ compares priors with observed co-function relationships. Co-function agreement and prior agreement were quantified as described above. Train–test similarity (*S*) was defined from an input similarity matrix and a given train–test split of perturbations. For each test perturbation, *S* was calculated as its average similarity to the *K* nearest training perturbations. *S*_cofunc_ was calculated using the co-function matrix, whereas *S*_prior_ was calculated by averaging *S* across five prior similarity matrices in the functional consensus group (GenePT, STRINGdb, MSigDB, DepMap, and scGPT). Stability with respect to *K* was assessed for *K* = 5, 10, and 20; *K* = 5 was used in the final implementation.

### Estimation of predictive performance for Perturb-seq datasets

To assess predictive performance of prior-informed models, we extended the benchmark framework of Miller et al. to six Perturb-seq datasets and an additional X-Atlas/Orion dataset [15–18,32]. Preparation and quality-control workflows described by Miller et al. were consistently applied across datasets. Briefly, perturbations were randomly selected with a maximum of 2,500s and selected transcription factors were added for gene regulatory context. For each model–dataset pair, models were trained using their original procedures or the corresponding CellSimBench adapters and evaluated by fivefold perturbation-level held-out cross-validation. For each held-out perturbation target, predictive performance was assessed using weighted *R*^2^(Δpert) between predicted and observed perturbation-induced expression changes for a fixed gene subset across held-out perturbations [32]. Results were then aggregated across folds and benchmarked against mean-predictor baselines.

For the retrospective training-set selection analysis, we randomly selected five test sets of 100 perturbations each using K562 genome-wide screen dataset. Within each repeat, the test set was held fixed while high- and low-similarity training sets were selected from the candidate perturbation pool at each evaluated training-set size. The two training-set conditions were matched on the distribution of *A*_prior_ and differed in prior similarity to the test set. The predictive model was trained separately on each selected training set and evaluated on the same test targets using Pearson(Δctrl).

## Declarations

### Ethics approval and consent to participate

Not applicable

## Consent for publication

Not applicable

## Availability of data and materials

All Perturb-seq datasets analyzed in this study were obtained from publicly available sources: the Genome-Wide Perturb-seq portal for Replogle et al. (2022) (https://gwps.wi.mit.edu/), the Gene Expression Omnibus under accession GSE264667 for Nadig et al. (2025) (https://www.ncbi.nlm.nih.gov/geo/query/acc.cgi?acc=GSE264667), and the X-Atlas/Orion dataset (https://huggingface.co/datasets/Xaira-Therapeutics/X-Atlas-Orion; https://doi.org/10.1101/2025.06.11.659105). The preprocessed matrices generated by the PRESAGE study were retrieved from the following Zenodo record (https://zenodo.org/records/15587986). Code for the analyses is publicly available at https://github.com/Yonsei-Hwang-Lab/page_reproducibility.

## Competing interests

The authors declare that they have no competing interests.

## Funding

This work was funded by grants from National Research Foundation of Korea (NRF) grants funded by the Korean government (MSIT; Grant Nos. RS-2025-17992868 and RS-2026-25488562). Also, We acknowledge the support of the Seok-San Biomedical Science Scholarship from Yonsei University College of Medicine.

## Author contributions

B.H. conceptualized this study.

J.P., H.S., and B.H. designed and performed data analysis.

J.P. and H.S. performed formal analysis.

J.P., H.S., and B.H. wrote the original draft.

## Acknowledgments

Not applicable

## Notes

### Competing Interest Statement

The authors have declared no competing interest.

