## Supplementary Materials for "Gene-level structure of prior knowledge and perturbation responses informs target-level predictability in prior-informed models"

**Supplementary Tables provided in a separate excel file**

**Supplementary Table 1. GSEA enrichment results on prior agreement (PC)**

**Supplementary Table 2. GSEA enrichment results on co-function agreement (FC)**

**Supplementary Table 3. GSEA enrichment results on co-regulation agreement (RC)**

**Supplementary Figures**

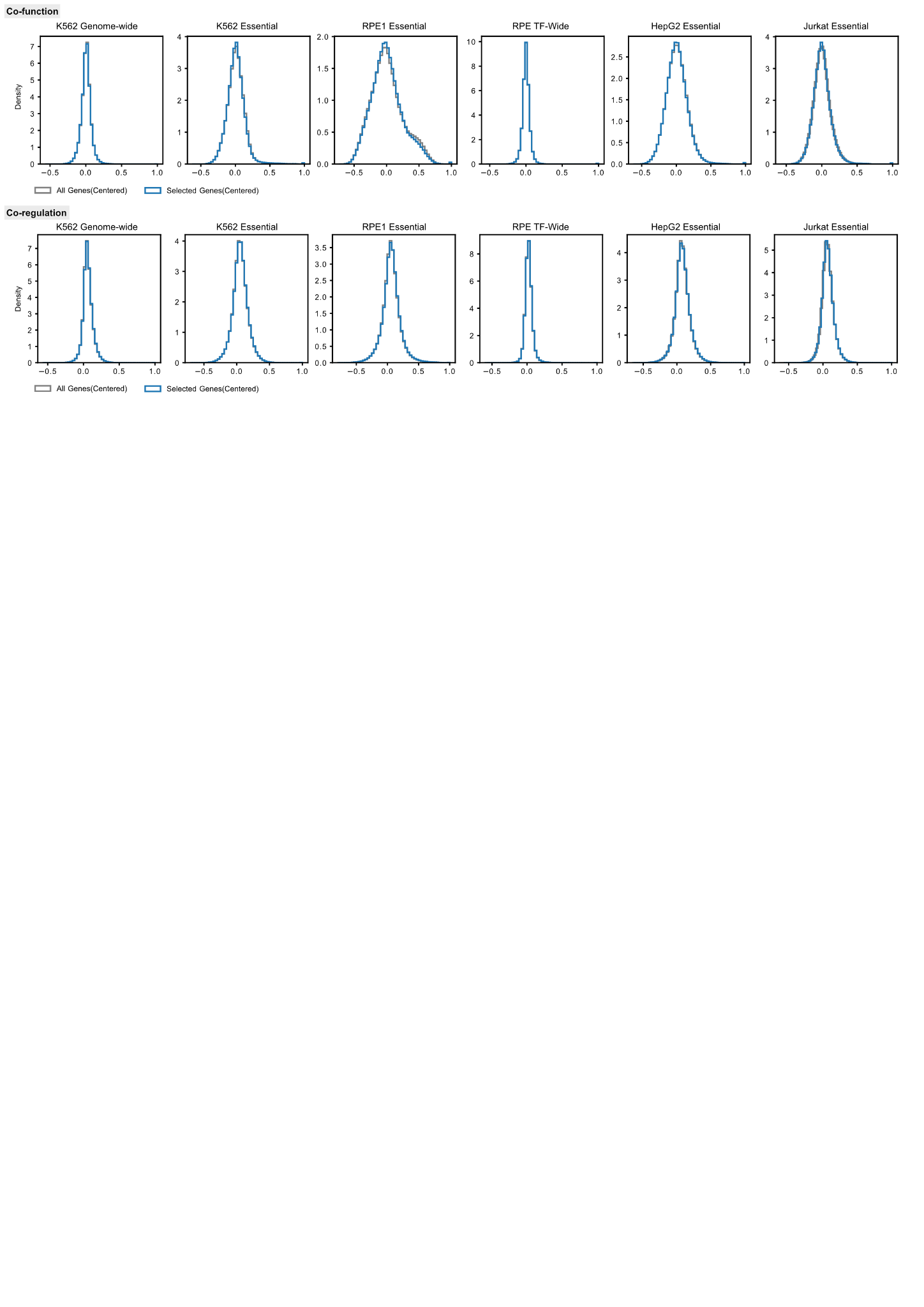

**Supplementary Figure 1. Effect of reference gene set subsetting on response-derived similarity.** Density distributions of co-function (top) and co-regulation (bottom) across six Perturb-seq datasets. Gray and blue traces show similarities calculated using all available genes and the subset in the reference gene set, respectively, after mean-centering perturbation responses.

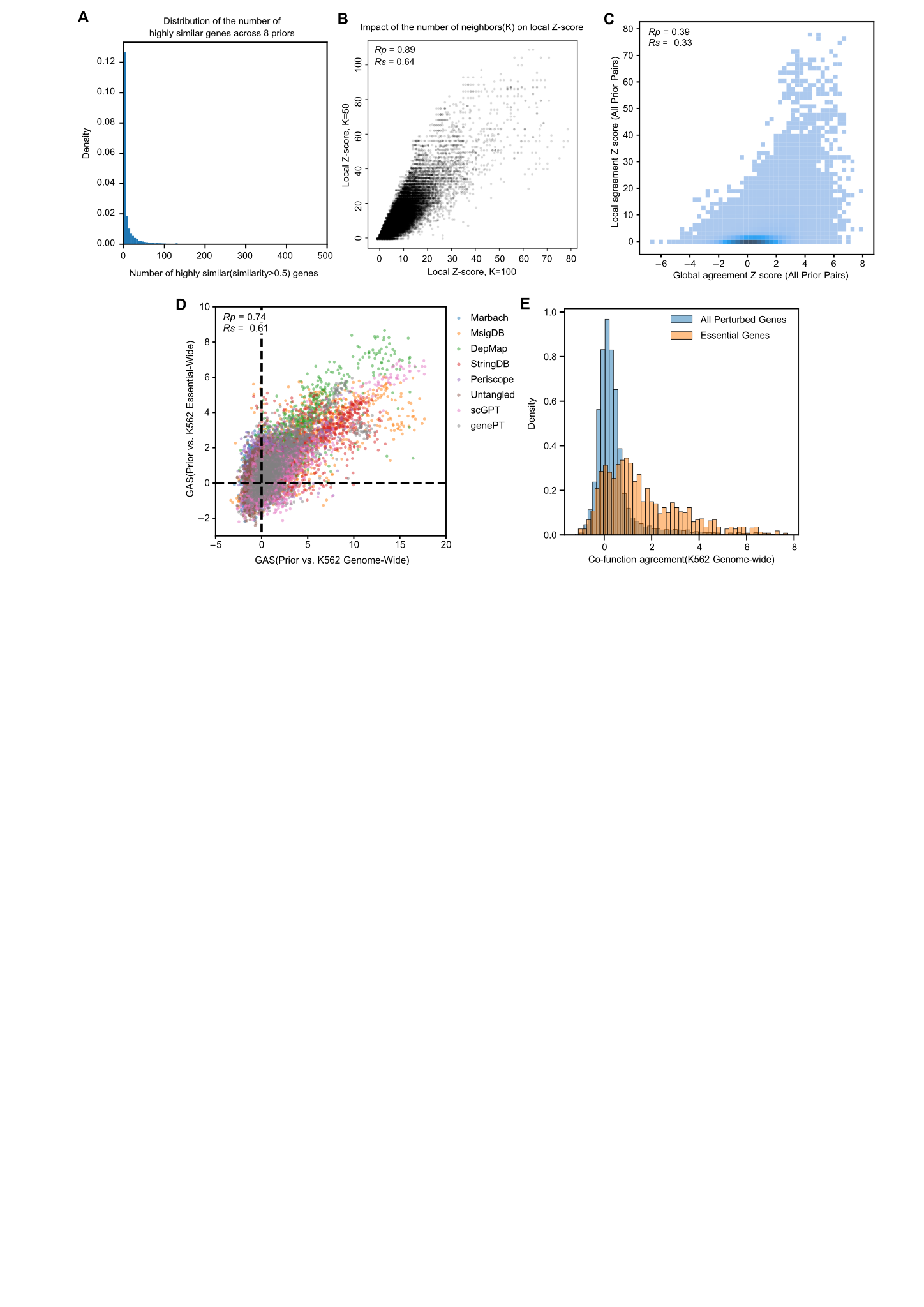

**Supplementary Figure 2. Stability and distribution of agreement measures.** (**A**) Distribution of the number of genes with similarity greater than 0.5 in each gene similarity profile, pooled across the eight prior knowledge sources. (**B**) Comparison of local-agreement Z-scores calculated using K = 50 and K = 100 nearest neighbors across all prior pairs. (**C**) Relationship between local and global Z-scores. (**D**) Concordance of GAS (gene agreement score) measured by comparing each prior with the K562 genome-wide screen (x axis) or the matched K562 essential-wide screen (y axis). (**E**) Distribution of co-function agreement in the K562 genome-wide screen for all perturbed genes and essential genes.

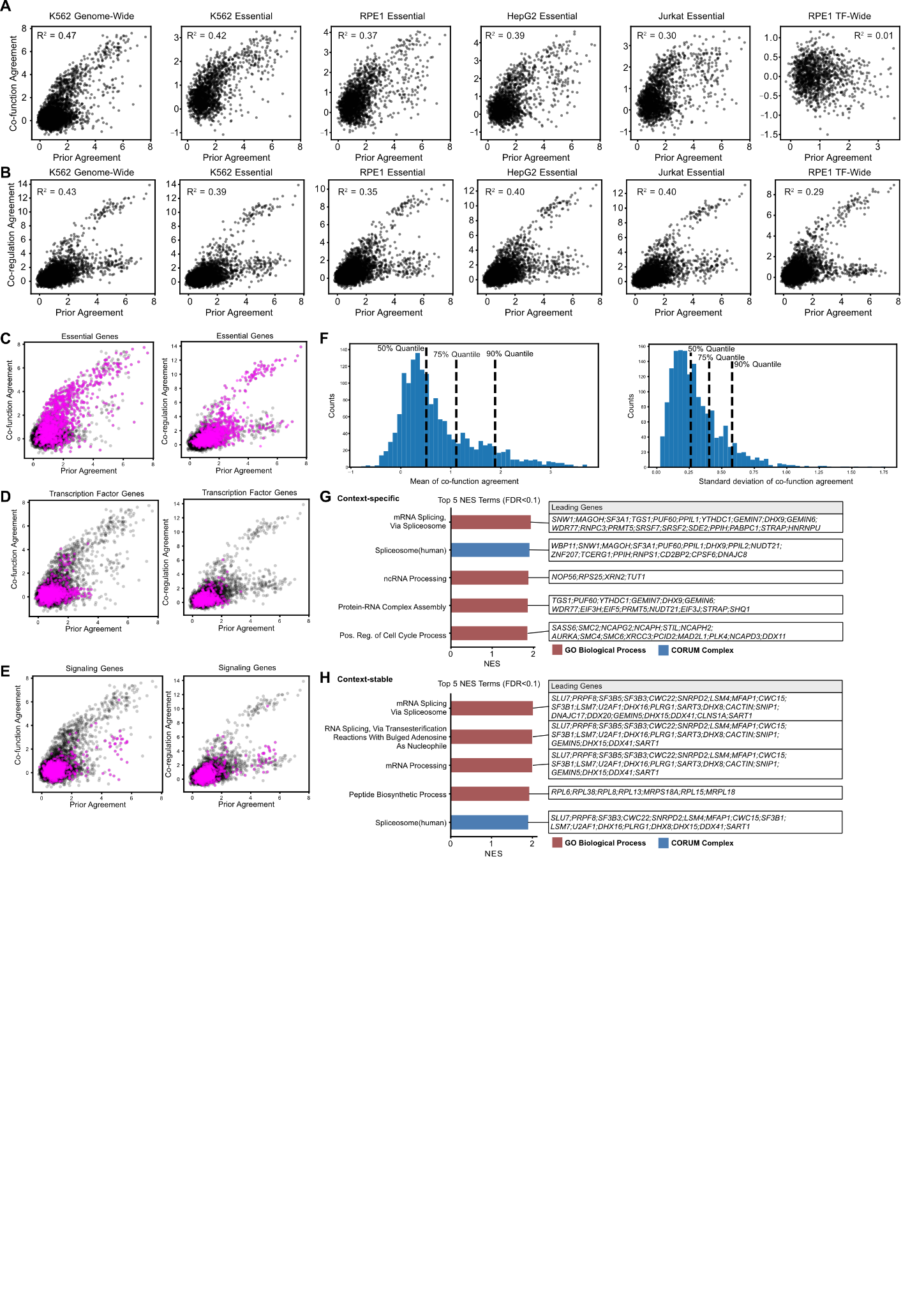

**Supplementary Figure 3. Agreement patterns across gene annotations and cellular context.** (**A, B**) Associations of prior agreement with co-function agreement (**A**) and co-regulation agreement (**B**) across six Perturb-seq datasets. R^2^ values indicate variance explained by prior agreement. (**C–E**) Prior agreement versus co-function (left) and co-regulation (right) agreement in the K562 genome-wide screen, with essential genes (**C**), transcription factors (**D**), and signaling genes (**E**) highlighted in magenta; other genes are shown in gray. (**F**) Distributions of the mean and standard deviation of co-function agreement across the four essential-wide screens for 1,625 shared perturbation-targets. Dashed lines indicate the 50th, 75th, and 90th percentiles. (**G, H**) Gene set enrichment analysis of context-specific (**G**) and context-stable (**H**) agreement patterns. Bars show the five significant terms with the highest normalized enrichment scores (FDR <` 0.1), and boxes list leading-edge genes. Colors denote GO Biological Process and CORUM Complex terms.

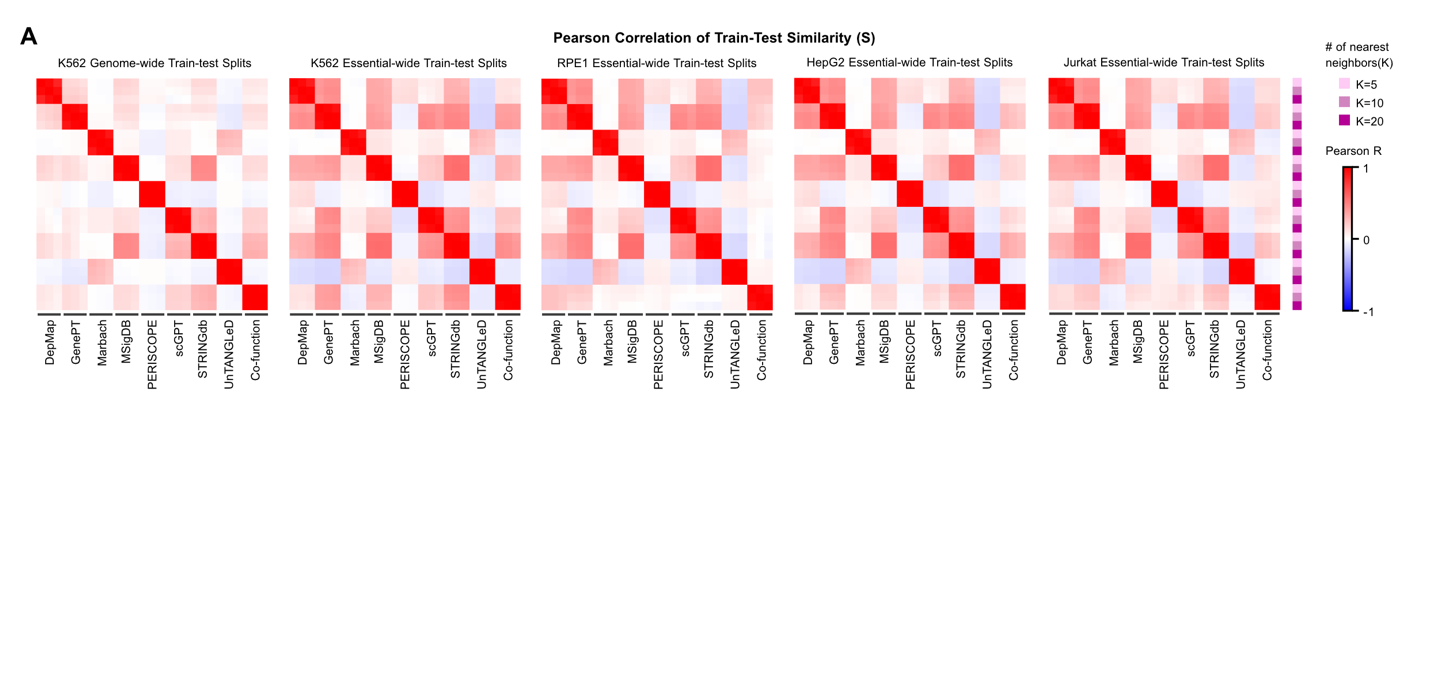

**Supplementary Figure 4. Stability of train–test similarity.** (**A**) Sensitivity of train–test similarity (*S*) to the number of nearest neighbors (K = 5, 10, or 20). Pearson correlations of train–test similarity across test perturbations, aggregated over five folds, for each similarity profile (prior knowledge or co-function) and value of K.

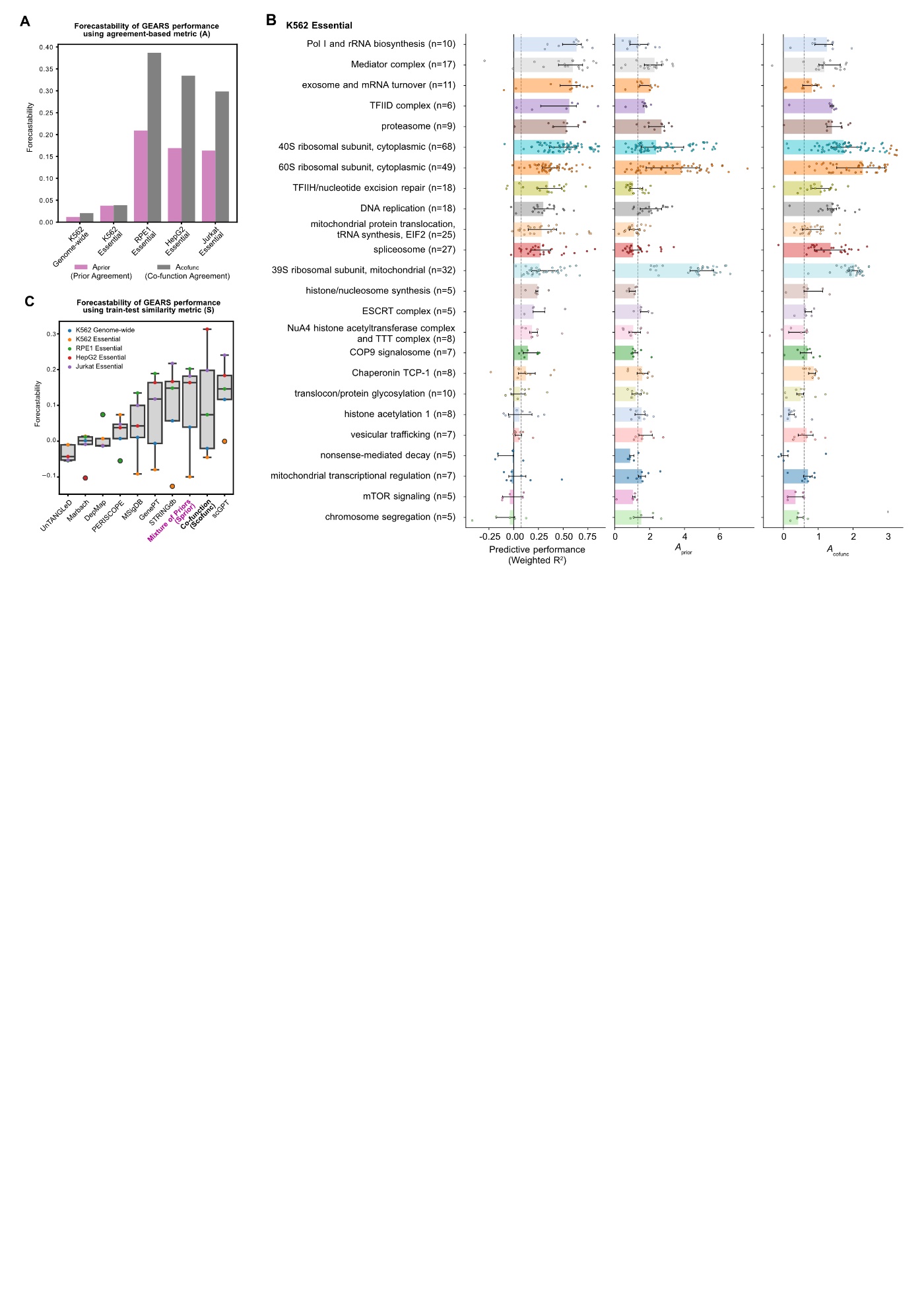

**Supplementary Figure 5. Gene-set level patterns of predictive performance and agreement, and forecastability of GEARS performance.** (**A**) Forecastability of GEARS performance using agreement-based metrics (*A*). (**B**) Predictive performance of PRESAGE (weighted *R*^2^), prior agreement (*A*_prior_), and co-function agreement (*A*_cofunc_) for genes grouped into functional gene sets in the K562 essential-wide screen. (**C**) Forecastability of GEARS performance using train–test similarity metrics (*S*) across individual priors, the five-prior average, and co-function.

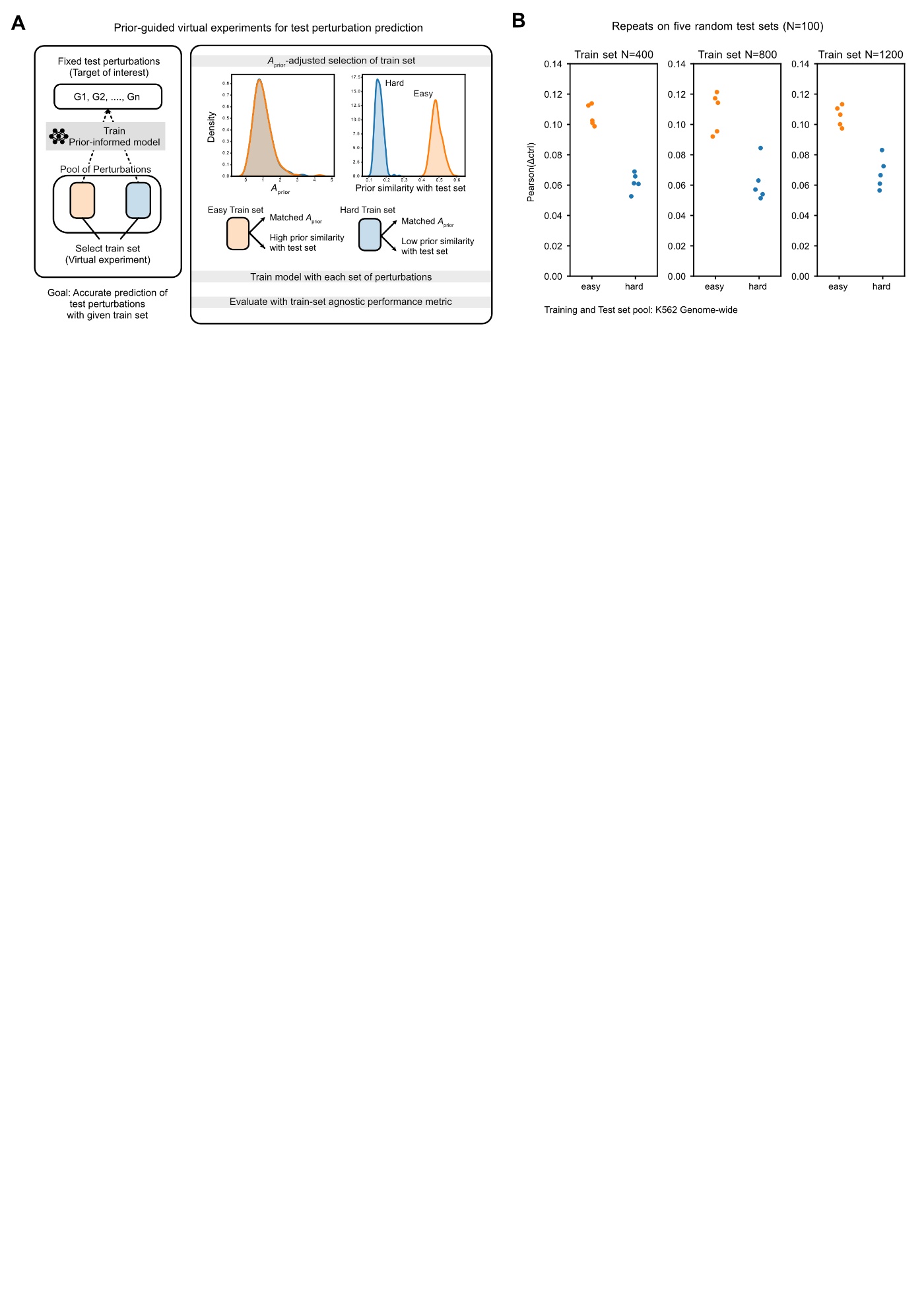

**Supplementary Figure 6. Prior-guided training-set selection for fixed test perturbations.** (A) Retrospective training-set selection design. High-similarity (Easy, orange) and low-similarity (Hard, blue) training sets have matched distributions of prior agreement (A_prior) and differ in their prior similarity to the same test targets. (B) Predictive performance, measured by Pearson(Δctrl), at the indicated training-set sizes. Each point represents one test-set repeat (five repeats; 100 randomly selected test perturbations per repeat). The same test targets are used for both training-set conditions within each repeat.
